# Akt/Foxo pathway activation switches apoptosis to senescence in short telomere zebrafish

**DOI:** 10.1101/2020.01.10.901603

**Authors:** Mounir El-Maï, Marta Marzullo, Inês Pimenta de Castro, Miguel Godinho Ferreira

**Author notes:** These authors contributed equally in this work (Mounir El-Maï, Marta Marzullo, Inês Pimenta de Castro).

## Abstract

Progressive telomere shortening during lifespan is associated with increased genome instability, block to cell proliferation and aging. Apoptosis and senescence are the two main cellular outcomes upon irreversible cell damage. In this study, we show a transition between apoptosis to senescence in cells of two independent tissues in telomerase zebrafish mutants. In young mutants, proliferative tissues exhibit defects in cell proliferation and p53-dependent apoptosis, but no senescence. Progressively, these tissues display signs of tissue dysfunction, loss of cellularity and increased senescence. These alterations are accompanied by an activation of pro-proliferative stimulus mediated by AKT. Consequently, FoxO1 and FoxO4 transcriptional factors are inactivated, reducing SOD2 levels, causing an increase in ROS. These alterations elicit the activation of the zebrafish p16/15 and senescence. Thus, upon telomere shortening in aging, early apoptosis induces compensatory proliferation. However, progressive decline in cell proliferation results in tissue damage and proliferative signals, promoting a switch to senescence.

## INTRODUCTION

Accumulation of DNA damage impairs cellular function and has been related to defects in tissue function, diseases and aging (Jackson and Bartek, 2009). To contrast the accumulation of damage, cells evolved DNA repair mechanisms. However when the damage persist cells undergo cell-cycle arrest, resulting in apoptosis or senescence (Childs et al., 2014).

Apoptosis is a programmed cell death, as a consequence of cell defects, and is highly regulated and p53-dependent. Apoptotic cells are usually eliminated from the tissue and replaced in highly proliferative tissues to maintain tissue homeostasis (Fogarty and Bergmann, 2017).

Senescence is a permanent cell-cycle arrest, generally associated with pro-inflammatory phenotype, known as Senescence Associated Secretory Phenotype (SASP) (Coppé et al., 2008). Senescent cells accumulates over time, and has been proposed that persistent SASP production can be associated to aging and age-related phenotypes (Krishnamurthy et al., 2004).

The CDK inhibitor (CKi) p16 encoded by the INK4A/ARF locus is a tumor suppressor that limits cell proliferation and is associated with cell senescence (Liao and Hung, 2003). It belongs to the INK family of CKIs that includes p15INK4b, p18INK4c and p19INK4d (Kamb, 1995; Vidal and Koff, 2000). The comparisons of the INK4a/ARF gene structure between man, mouse, chicken and the fugu fish revealed a dynamic evolutionary history of this locus (Gilley and Fried, 2001; Kim et al., 2003). p15INK4b (p15), a close relative of p16, is encoded by the INK4b gene, located immediately upstream of the INK4a/ARF locus. Similar to p16, this CKI has been associated to cell senescence (Fuxe et al., 2000; Hitomi et al., 2007; Senturk et al., 2010). In chicken, as in fugu fish, INK4a is mutated and does not encode a functional protein. Thus, fugu fish INK4a locus expresses Arf but not p16 (Liao and Hung, 2003), suggesting that the function of p16 was possibly being taken over by p15.

Under physiological conditions, cells with a high turn-over rate, including epithelial and germinal cells, preserve tissue homeostasis by undergoing continuous cellular proliferation and apoptosis. Nevertheless, cells can only go through a finite number of divisions before entering an irreversible cell-cycle arrest, process known as replicative senescence. Telomere erosion has been proposed to constitute the “molecular clock” that determines the number of divisions a cell can undergo before reaching senescence, phenomenon known as Hayflick limit (Bodnar et al., 1998; Hayflick, 1965).

Telomeres are nucleoprotein complexes that protect the extremities of linear chromosomes and counterbalance incomplete replication of terminal DNA (Jain and Cooper, 2010; O’Sullivan and Karlseder, 2010). In most eukaryotes, the end replication problem is solved by telomerase, which expression is restricted in most human somatic cells (Forsyth et al., 2002). Consequently, telomeres shorten significantly during human aging (Aubert and Lansdorp, 2008).

Vertebrate telomerase mutant animal models have been used to assess the direct association between telomere shortening and tissues dysfunction. Late generation (G4-6) telomerase knockout mice present premature features of aging, including reduced cell proliferation and increased apoptosis of several tissues (Lee et al., 1998; Rudolph et al., 1999). In zebrafish, first generation telomerase mutants (tert−/−) present short telomeres and promptly display premature aging phenotypes including tissue decline (Anchelin et al., 2013; Carneiro et al., 2016a; Henriques et al., 2013).

In the absence of telomerase, telomeres become critically short, accumulate γH2A.X and activate the DNA Damage Response (DDR) (d’Adda di Fagagna et al., 2003). One of the mediators of DDR is the onco-suppressor p53, which accumulates upon telomere shortening and causes either cell senescence or apoptosis (Li et al., 2016). So far, it is unclear what are the signals determining one or the other cell fate in response to p53 accumulation.

Previous studies suggested that cellular senescence is associated with increased levels of mTORC/AKT signaling (Miyauchi et al., 2004; Moral et al., 2009). AKT is a serine/threonine protein kinase that is activated upon pro-proliferative extracellular signals. This pathway is triggered by growth factor receptors, including the Insulin Growth Factor Receptor (IGFR) (Liao and Hung, 2010). Activation of AKT and mTORC2-mediated phosphorylation result in the phosphorylation of the forkhead transcriptional factors, FoXO1 and FoxO4 (Tuteja and Kaestner, 2007). Once phosphorylated, the FoXO family proteins translocate outside the nucleus thus resulting in the repression of their main target genes, including the superoxide dismutase SOD2. Prolonged and uncontrolled activation of this pathway results in mitochondrial dysfunction and increased ROS levels (Nogueira et al., 2008).

In this study, we investigated the in *vivo* switch between apoptosis and cell senescence as a consequence of telomere shortening in tert−/− zebrafish. We describe that early in life, telomerase deficiency results in p53 mediated apoptosis and loss tissue homeostasis, causing pro-proliferative AKT pathway activation in older individuals. AKT/FoxO signal cascade then triggers a switch that results in mitochondrial dysfunction, increased levels of ROS and p15/16 leading to cell senescence.

## RESULTS

### *tert−/−* zebrafish proliferative tissues undergo a time-dependent switch from apoptosis to cell senescence

Apoptosis is a process in which programmed cell death allows for clearance of damaged cells (Hawkins and Devitt, 2013). In contrast, replicative senescence is a state of terminal proliferation arrest, associated to gradual telomeres attrition occurring during cell division (Olovnikov, 1973; Shay and Wright, 2000). To explore the molecular mechanisms underlying the cell-fate decision between apoptosis and senescence, we used telomere attrition as a trigger of these two possible outcomes.

First-generation tert−/− zebrafish have shorter telomeres than their wild-type (WT) siblings, develop several degenerative conditions affecting mainly highly proliferative tissues, such as the testis and gut, and die prematurely (Anchelin et al., 2013; Carneiro et al., 2016a; Henriques et al., 2013). At 3 months of age, tert−/− fish are macroscopically similar to their WT siblings (Carneiro et al., 2016a), with testis and gut being histologically indistinguishable from WT (Figure 1A). However, at this early age, average telomere length is short and trigger the onset of DDR and increased apoptosis in tert mutants (Carneiro et al., 2016a). We analyzed the presence of apoptotic cells in 3 months-old tert−/− gut and testis using the TUNEL assay. We confirmed that, even in the absence of macroscopic defects, tert−/− gut and testis exhibits a higher number of apoptotic TUNEL-positive cells compared to their WT siblings (Figure 1C). In order to confirm the activation of DDR, we analyzed the phosphorylation levels of the DNA damage marker γH2A.X on whole cell lysates from gut and testis of 3 month-old tert−/− zebrafish (Figure 1E, quantification in Supp. Figure 1). We detected a significant increase of the ATM-dependent phosphorylated form of γH2A.X in ser139 in both 3 month-old tert−/− gut and testis (Figure 1E, quantification in Supp. Figure 1). As expected, we observed a concomitant increase p53 protein levels in both tissues of tert−/− zebrafish (Figure 1E, quantification in Supp. Figure 1).

**Figure 1.**
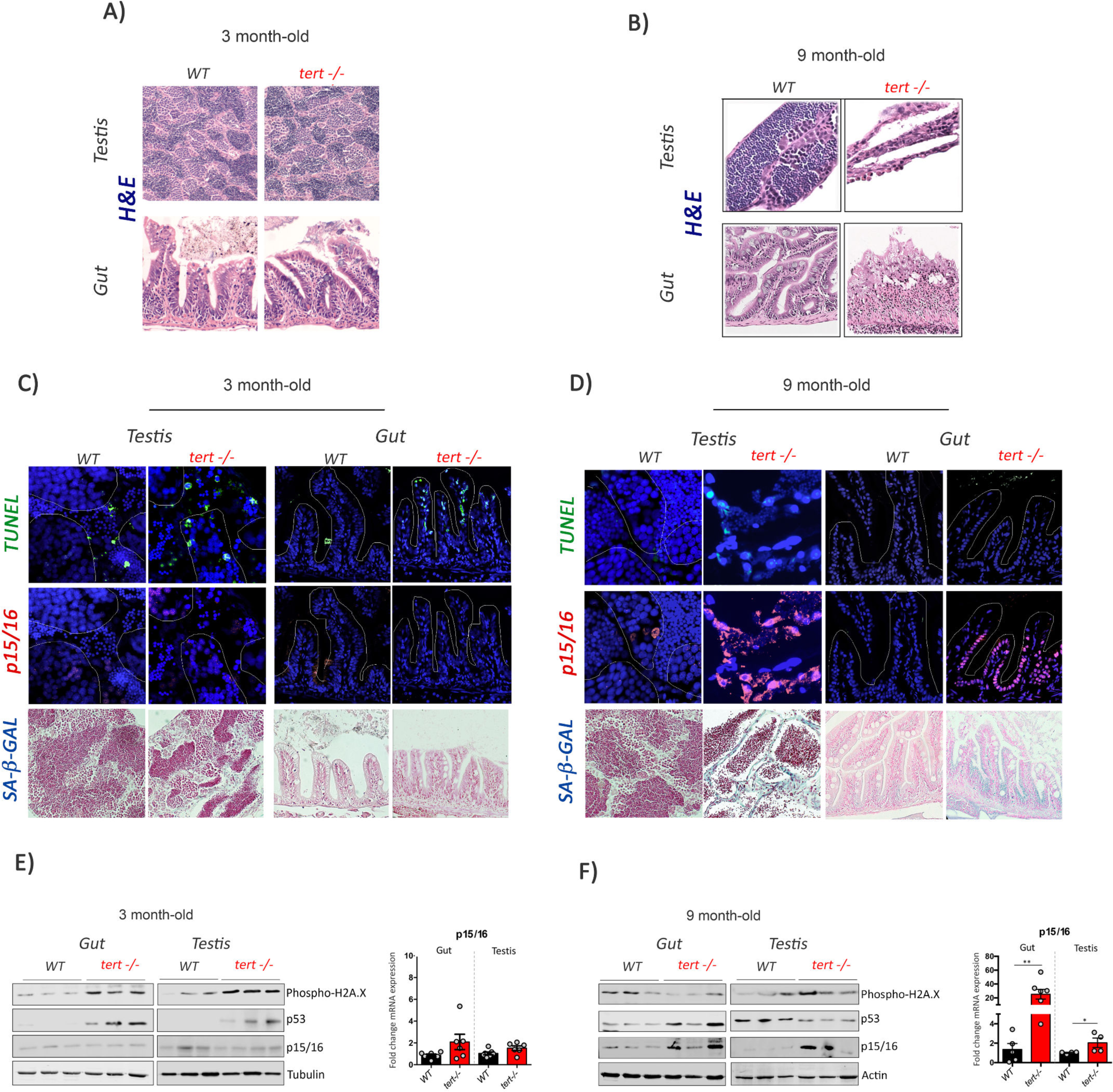
Proliferative tissues of tert−/− zebrafish exhibit a switch from apoptosis to senescence with age. A-B) Representative haematoxylin and eosin-stained sections of gut and testis from 3 month (A) or 9 month-old (B) WT and tert−/− siblings. While no macroscopic tissue defects are distinguishable at 3 months (N=3), 9 month-old tert mutants (N=3) exhibit altered gut and testis structures. C-D) Representative immunofluorescence images of apoptosis (TUNEL) or senescence (p15/p16 or SA-β-GAL) of gut and testis from 3 month (C) or 9 month-old (D) WT and tert−/− siblings (N=3 each). Dashed outlines locate zones of maturing spermocytes (testis) or villi (gut). At 3month, both tissues show an increased number of apoptotic cells in tert−/− compared to WT. At that age, no signs of senescence are visible in these tissues. However, senescent cells appear in gut and testis of 9 month-old tert−/− fish depicting a switch between apoptosis and senescence at that age. E-F) Western blot and RT-qPCR analysis for DNA Damage and senescence associated genes in gut and testis of 3 month (E) or 9 month-old (F) WT and tert−/− siblings (N>=6 fish). RT-qPCR graphs are representing mean ±SEM mRNA fold increase after normalisation by RPL13a gene expression levels (* p-value <0.05; ** p-value<0.01). At 3 month, gut and testis showed higher levels gH2AX and p53 proteins in tert−/− compared to WT but no differences in P15/p16 expression. At 9 month, senescence-associated p15/16 expression increased in tert mutant compared to WT siblings.

In light of the differences found between the INK4a/ARF locus in zebrafish and mammals, we decided to test the conservation of the protein and the validity of the mammalian anti-p16 antibody (sc-1661, Santa Cruz Biotechnology) used for the senescence analysis. To this purpose, we designed antisense morpholino oligonucleotides (p15/16 MOs) and injected increasing amounts in 1 cell-stage embryos (Supp. Figure 2). A control morpholino sequence was included as negative control (CTR MO). At 3dpf upon morpholino injection, larvae were collected and tested for the expression of the p15/16 protein in zebrafish. Western Blot analysis revealed that injection of increasing concentration of p15/16 MOs causes a reduction in the amount of the protein recognized by the anti-p16 antibody, indicating that the sequence of the protein associated to senescence is conserved from mammals to zebrafish (Supp. Figure 2).

**Figure 2.**
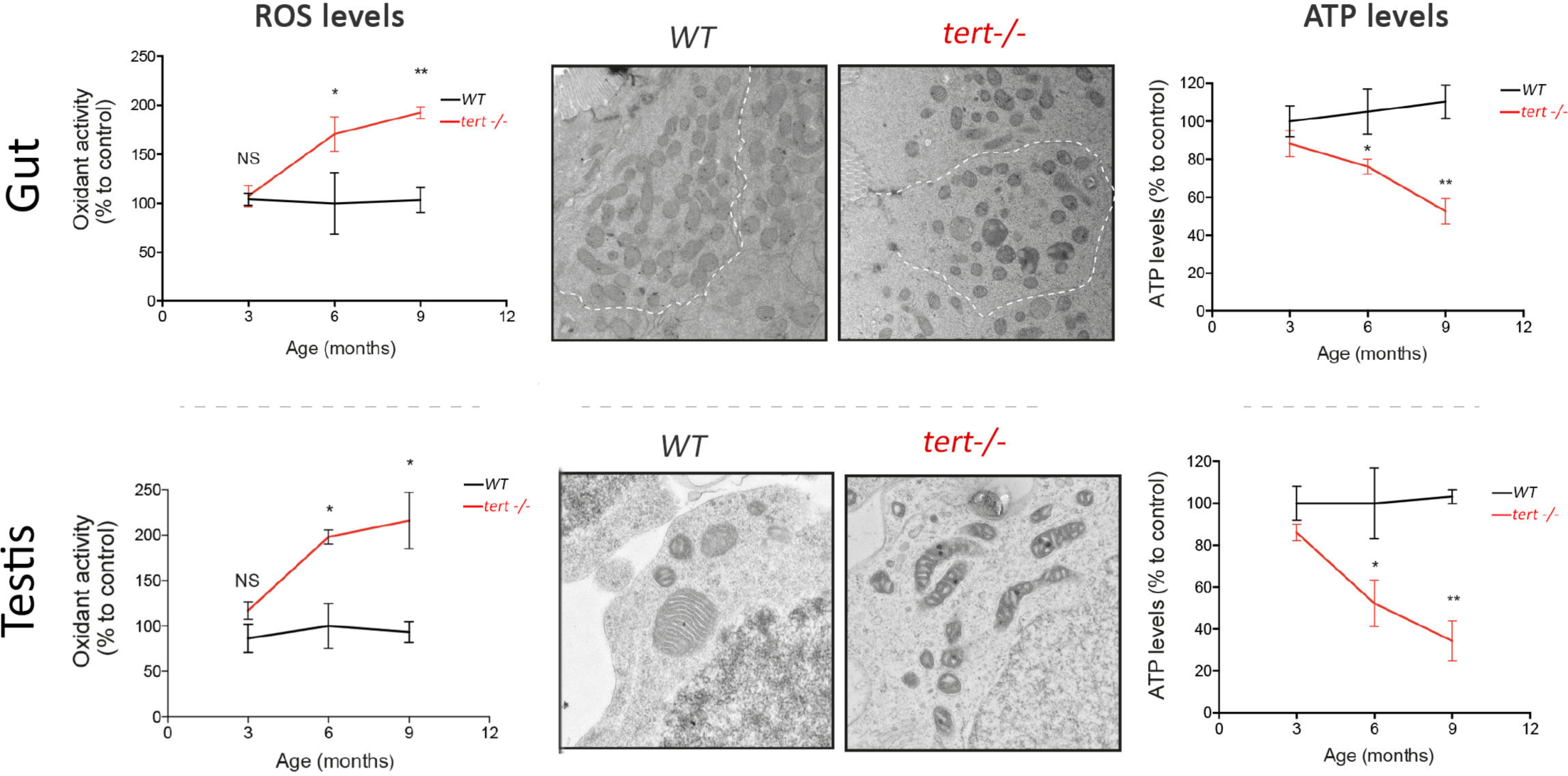
Gut and testis of tert−/− zebrafish are characterised by a time-dependant mitochondrial defect, increase of ROS levels and reduction of ATP levels. Compared to WT where difference are seen from 3 to 9 month, gut and testis of tert−/− exhibit a time dependant increase in ROS level and decrease of ATP levels (N>=3 fish per time point per genotype). Representative EM images of these tissues at 9 month revealed fragmented mitochondrial ultrastructure in tert−/− testis and rounded and swollen mitochondria containing perturbed crystal structures (N>=3 fish). Data are represented as mean±SEM.

Strikingly, even though DDR is active in 3-month-old tert−/−, the analyzed tissues did not exhibit signs of cellular senescence. We were unable to detect senescence-associated beta-galactosidase (SA-beta-Gal) activity in both gut and testis in tert−/− fish (Figure 1C). Accordingly, we observed no differences in expression of the senescence marker p15/16 by qRT-PCR, western blot (Figure 1E) or by immunofluorescence staining (Figure 1C). Therefore, at 3 month tert−/− in which tissue integrity is retained, telomere dependent DDR signalling predominantly induces apoptosis but no detectable cell senescence.

To investigate the consequences of telomere erosion and chronic DDR activation in aging, we analysed testis and gut of older tert−/− animals (9 month of age). Contrary to what we observed in 3 month old fish, older tert−/− zebrafish exhibit tissue morphological defects (Figure 1B), including testis atrophy and width lengthening of the gut lamina propria (as described previously -(Carneiro et al., 2016a)-).

Because telomere shortening is known to induce both apoptosis and cell senescence, we wondered if the decline in tissue homeostasis represented a change in cell fate. Surprisingly, at 9 month of age, we could not observe clear differences in p53 levels between WT and tert−/− (Figure 1F). In fact, tert−/− gut and testis exhibited a decline in apoptosis in 9 month-compared to 3 month-old fish, denoting a decrease in TUNEL positive cells (Figure 1D). In contrast, at this stage, these tissues exhibited a clear accumulation of senescent cells in tert−/− compared to WT, as revealed by SA-beta-Gal staining (Figure 1D). Increased senescence was confirmed by an increase in p15/p16 by immunofluorescence (Figure 1D), mRNA and protein levels (Figure 1E). In addition, we observed that reduction of apoptotic cells and increase of senescent cells was concomitant with higher levels of expression of Bcl-XL mRNA suggesting an activation of anti-apoptotic pathways in old tert−/− fish (Supp. Figure 3). Taken together, these results show in vivo a switch from apoptosis to senescence during aging of tert mutant fish, and that this switch associates with age-dependent tissue degeneration.

**Figure 3.**
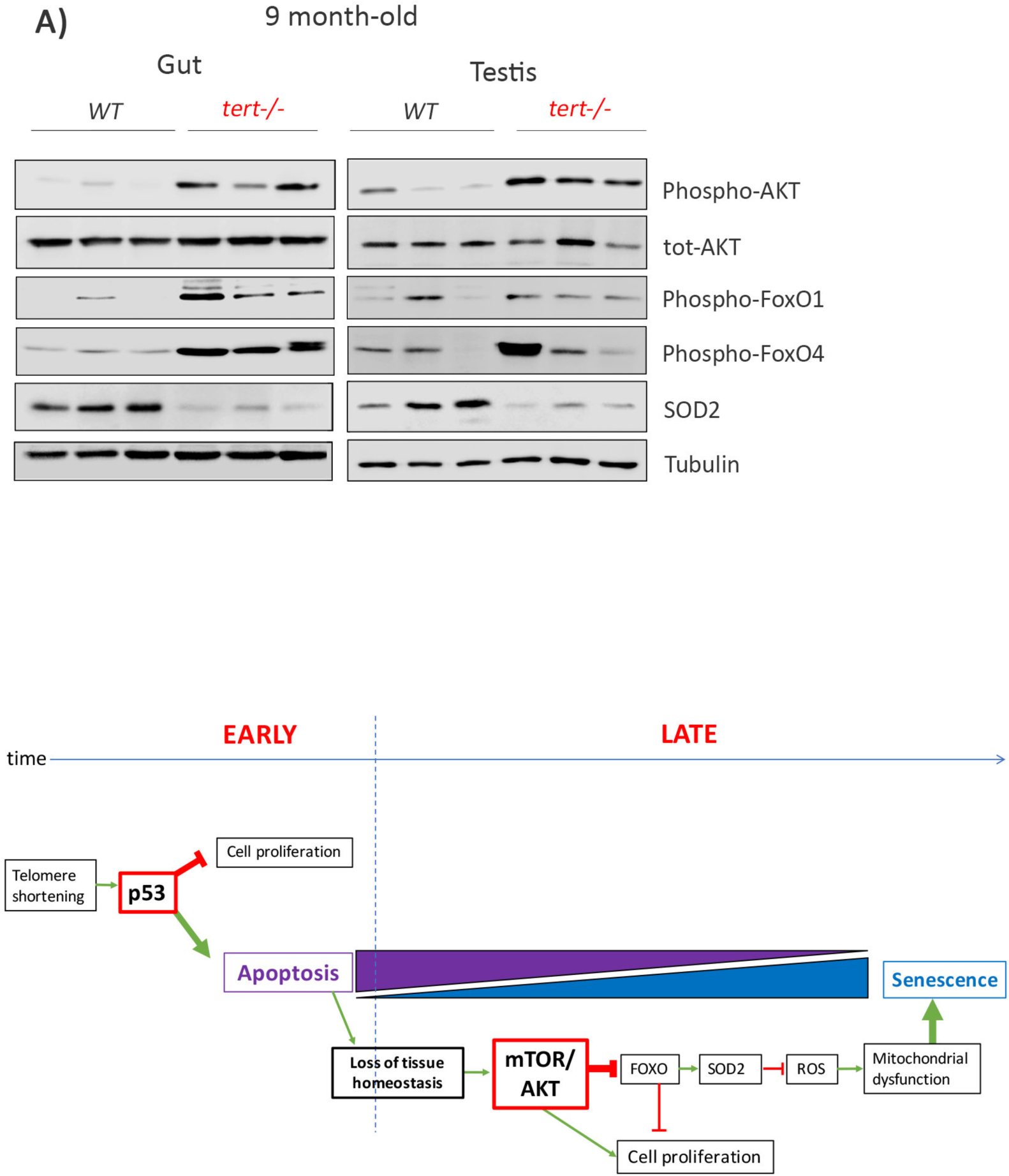
Activation of Akt in old tert−/− leads to ROS accumulation by blocking the FoxO1/4-SOD2 Axis and promoting mitochondrial dysfunction. Representative immunoblot of p-Akt, total Akt, pFOXO1, pFOXO4 and SOD2 from testis and gut of 9 month-old tert mutant and WT siblings (N>=9). At 9 month, these proliferative tissues show an increased activation of Akt leading to the inhibition of FOXO-dependant SOD2 expression.

### ROS accumulation and mitochondrial dysfunction become apparent upon short telomere-induced senescence

In mammalian systems, similarly to what we observe in zebrafish, DNA damage initially halts cell-cycle progression through a p53/p21-mediated cell-cycle arrest (Rodriguez and Meuth, 2006) (Figure 1E). But if lesions persist, expression of p16Ink4a predominates as a consequence of mitochondrial dysfunction and ROS production (Freund et al., 2011; Passos et al., 2010). Late generation telomerase knockout mice were observed to induce mitochondrial dysfunction through p53-dependent suppression of the master regulator of mitochondrial biogenesis, PGC1α (Sahin et al., 2011). G4 mTERT deficient mice exhibit significant alterations in mitochondrial morphology, accumulation of ROS and reduced ATP generation (Sahin et al., 2011).

We investigated if mitochondrial dysfunction could play a part in the apoptosis to senescence switch observed in tert−/− zebrafish. First, we started by examining if p53 activation triggers the repression of PGC1α in zebrafish. Curiously, despite significant accumulation of p15/16 and p53 (Figure 1F), we did not observe differences in terms of RNA or protein levels of PGC1α in older tert−/− gut extracts (Supp. Figure 4). However, we did detect a robust increase in oxidative damage with age. By 3 months of age, the levels of ROS in tert−/− gut and testis do not differ significantly from their WT siblings (Figure 2A). Later, we observed a gradual and significant accumulation of ROS in both tissues from 6 months onward in tert−/− compared to WT controls (Figure 2A). Production of ROS, especially superoxide, is a necessary by-product of mitochondrial respiration (Murphy, 2009). Mitochondrial dysfunction is characterized by concurrent high superoxide production leading to a breakdown of membrane potential that compromises energy production and cellular metabolism (Balaban et al., 2005). In agreement with previous findings, we observed that testis mitochondrial ultrastructure became significantly fragmented in older tert−/− zebrafish (arrows, Figure 2B). Similarly, gut mitochondrial morphology became increasingly rounded and swollen with the appearance of perturbed crystal structure (arrows, Figure 2B). Consistently, we observed a significant reduction of levels of ATP in both tissues of tert mutants (Figure 2C). Together, these results indicate that mitochondrial function declines dramatically during aging of tert−/− proliferative tissues, supporting the idea that a change in mitochondrial homeostasis may dictate the tissue’s cell-fate decision.

**Figure 4.**
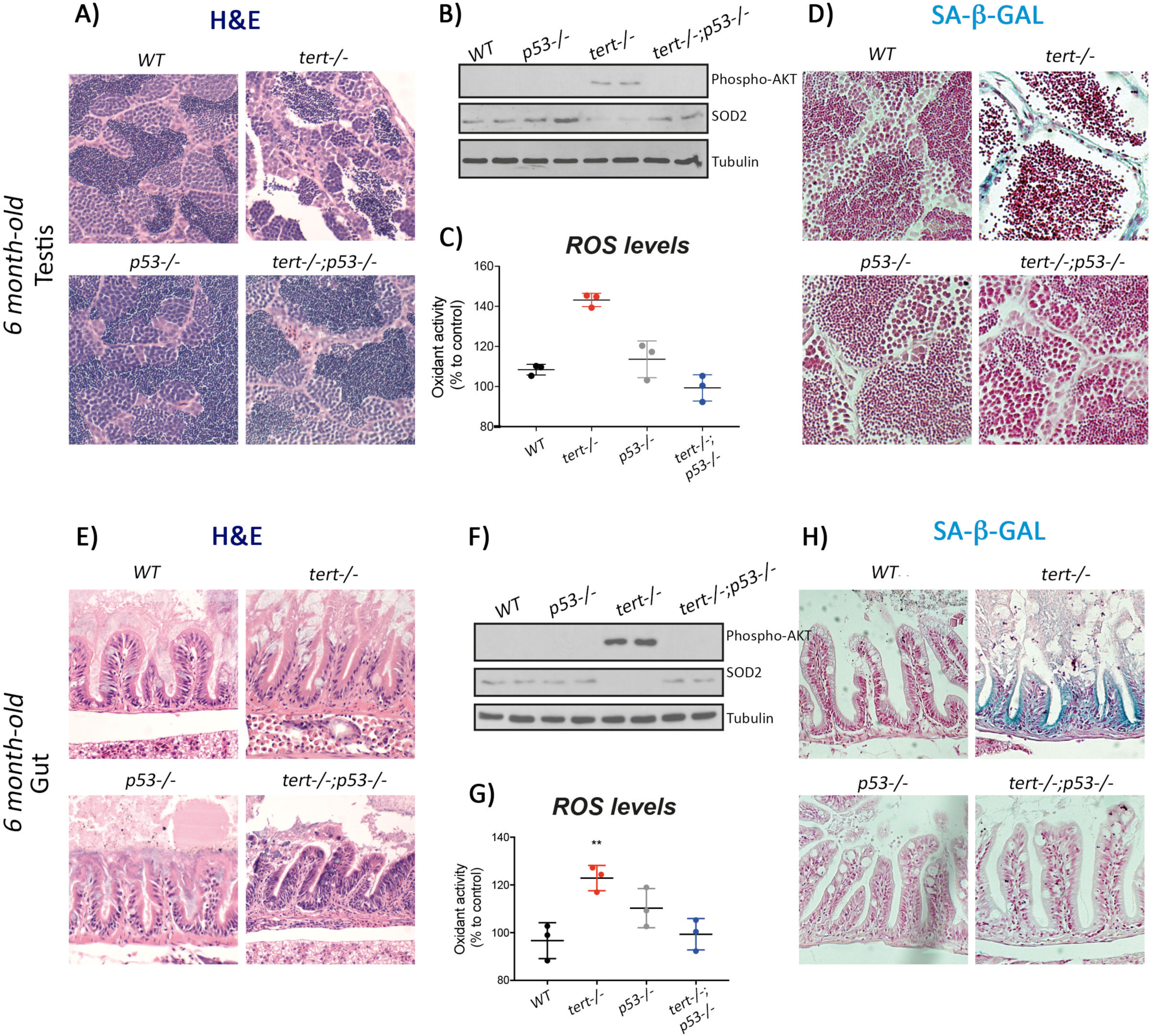
Genetic inhibition of p53 prevents short telomeres-induced tissue degeneration, Akt activation, ROS accumulation and induction of senescence. A and E) Representative haematoxylin and eosin-stained sections of testis (A) and gut (E) from 6 month-old WT, tert−/−, p53−/− and tert−/−/p53−/− siblings (N=3 fish each). Genetic inhibition of p53 rescues short-telomere dependant morphological defects of these tissues. B and F) Representative western blot analysis of pAkt and SOD2 in testis (B) and gut (F) (N=2 fish each). Inhibition of p53 impedes phosphorylation of Akt and downstream downregulation of SOD2 leading to a rescue of increased ROS levels (C and G; N=3 fish per genotype) seen in tert−/−. D and H) Representative images of SA-β-GAL staining of testis (D) and gut (H) from 6 month-old WT, tert−/−, p53−/− and tert−/−;p53−/− siblings (N=3 fish). Inhibition of p53 counteract telomere shortening-induced senescence.

### AKT promotes ROS production by blocking the FoxO1/FoxO4-SOD2 molecular axis

Excessive ROS formation gives rise to oxidative stress, leading to cellular damage and, eventually, senescence (Velarde et al., 2012). Mitochondrial manganese superoxide dismutase (SOD2) is one of the major ROS scavengers. Notably, SOD2 expression decreases with age (Tatone et al., 2006; Velarde et al., 2012). SOD2 KO mice and connective tissue-specific SOD2 KOs have reduced lifespan and exhibit premature aging phenotypes associated with senescence but no onset of apoptosis (Treiber et al., 2011; Velarde et al., 2012).

To gain mechanistic insights into the nature of the oxidative damage observed in the tert−/− zebrafish, we decided to analyse the expression levels of this important antioxidant defence enzyme. Western blot analysis of 9 months-old testis and gut samples, showed a significant decrease in terms of protein levels of SOD2 in tert−/− mutants compared to WT (Figure 3A). In contrast, SOD2 levels were not affected in tert−/− at 3 months of age (Supp Fig. 5). This result suggests that the mechanism that copes with superoxide production is compromised in older tert−/− mutants and, therefore, possibly responsible for the accumulation of oxidative damage in the affected tissues.

**Figure 5.**
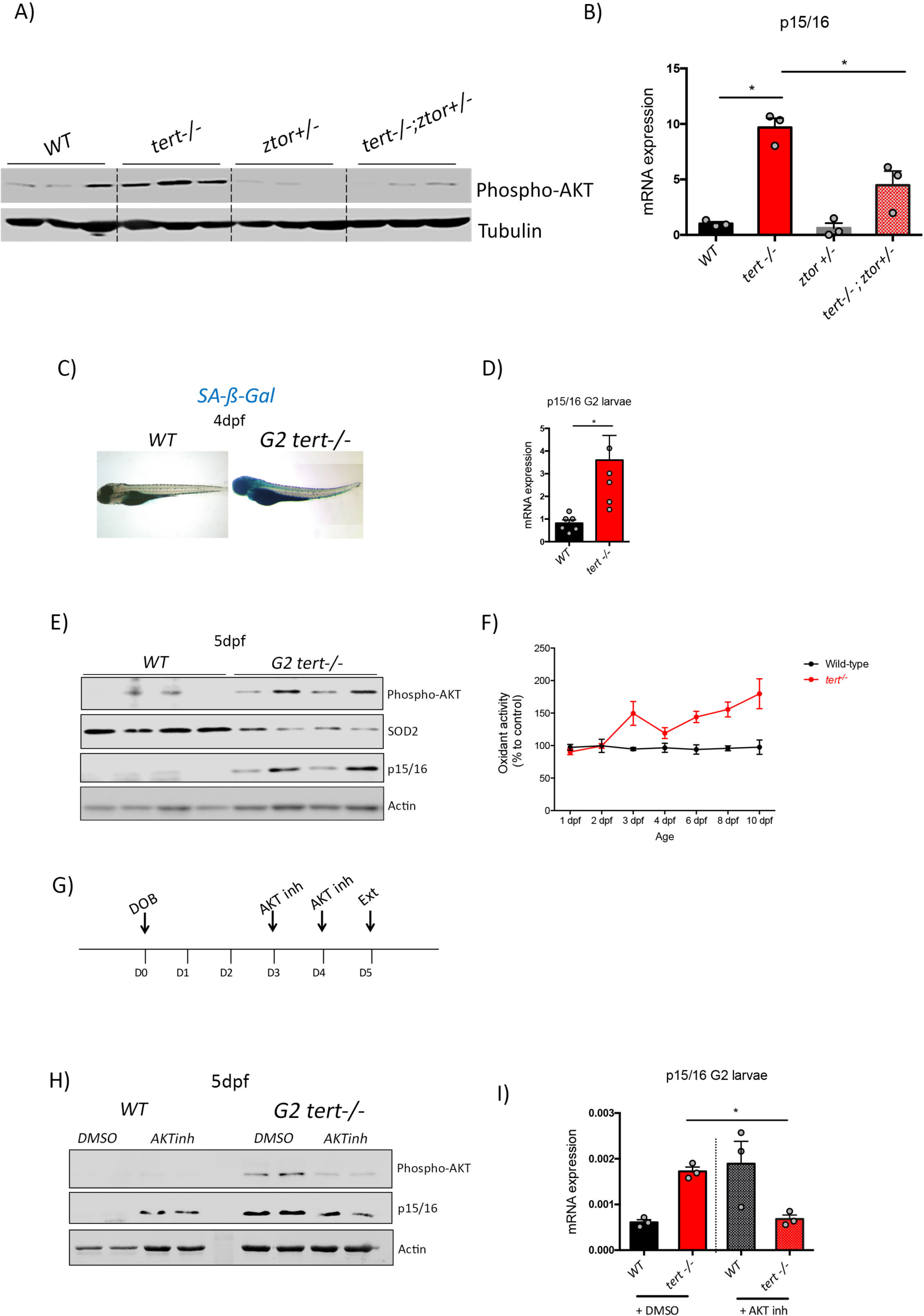
Genetic and pharmacological inhibition of Akt prevents short telomeres-induced senescence. A)Western blot analysis of pAkt and B) RT-qPCR analysis of p15/p16 mRNA levels in 11 month-old gut of WT, tert−/−, ztor+/− and tert−/−;ztor+/− fish (N=3 fish). Heterozygous mutation of zTOR counteracts telomere-shortening induced Akt activation leading to a rescue of p15/p16 induction. C) Representative images of SA-β-GAL staining, D) RT-qPCR analysis of p15/p16 mRNA levels (N=6), E) Western blot analysis of pAkt, SOD2, p15/p16 (N=4) and F) ROS levels measurements (N=3) in whole G2 tert mutant and WT larvae. Second generation of tert mutants exhibiting extremely short telomeres show premature aging phenotype described in Fig 1, 2 and 3 at larval stages. G) Experimental scheme of pharmacological inhibition of Akt in G2 tert−/−. H) Western blot analysis of pAkt and p15/p16 and I) RT-qPCR analysis of p15/p16 mRNA levels of G2 tert−/− and WT treated with Akt inhibitor. Pharmacological inhibition of Akt rescues telomere-shortening induced p15/16 expression. RT-qPCR graphs are representing mean ±SEM mRNA fold increase after normalisation by RPL13a gene expression levels (* p-value <0.05; ** p-value<0.01)

Phosphorylation (inactivation) of FoxO-family by the AKT kinase causes the elevation of intracellular ROS levels through the repression of detoxifying enzymes, such as SOD2 (Brunet et al., 1999; Kops et al., 2002; Miyamoto et al., 2007). FoxO proteins are a family of transcription factors responsible for a wide range of cellular processes, including cell cycle arrest, DNA damage response, metabolism and ROS detoxification (Greer and Brunet, 2005). Phosphorylation of FoxO by AKT triggers the rapid relocalization of FoxO from the nucleus to the cytoplasm, with the consequent downregulation of FoxO target genes. We, therefore, hypothesised that activation of AKT/FoxO signalling was responsible for the increased oxidative stress in older tert−/− zebrafish. As expected, increased phosphorylation levels of FoxO1 and FoxO4 were correlated with lower expression levels of SOD2 (Figure 3A, quantification in Supp. Figure 6). Phosphorylation of FoxO proteins suggests that inactivation of these transcription factors may be the cause for the down-regulation of SOD2 in older tert−/− gut and testis.

AKT is a highly conserved central regulator of growth-promoting signals in multiple cell types. The kinase activity and substrate selectivity of AKT are principally controlled by phosphorylation sites. Phosphorylation of serine 473 (pAKT-Ser473), is a consequence of activation of mammalian target of rapamycin complex 2 (mTORC2) (Sarbassov et al., 2005). pAKT-Ser473 is required for phosphorylation and inactivation of the FoxOs (Guertin et al., 2006). Accordingly, while we did not observe differences in total AKT protein levels, we detected a significant increase in the phosphorylated levels of pAKT-Ser473 denoting full activation of AKT in older but not younger tert−/− zebrafish (Figure 3A, quantification in Supp. Figure 6). Therefore, AKT activation correlates with increased levels of FoxO inhibitory phosphorylation and concomitant decrease in SOD2 protein levels in tert−/− mutants when compared to WT. Collectively, our results suggest that, upon tissue damage in older tert−/− zebrafish, activation of a pro-proliferative signaling pathway leads to AKT-dependent inactivation of FoxO1 and FoxO4, which in turn causes the down-regulation of SOD2 expression. These events impact on oxidative stress, triggering p15/p16 accumulation, causing a consequent senescent (irreversible) cell-cycle arrest.

### p53−/− rescue of tert−/− apoptosis delays the appearance of tissue degeneration and cellular senescence

Activation of a pro-proliferative pathway in an organism that exhibits defects in cell proliferation was somehow surprising. However, one major difference between 3 and 9 months-old gut and testis was the increasing tissue damage (Figure 1 A and B). In addition to our previous observation of reduction in the number of cell divisions (Carneiro et al., 2016a), tissue damage may be explained by increased cell death that predominates 3 months-old tert−/− testis and gut (Figure 1C).

In order to compensate for the empty space, dying cells in proliferative tissues induce compensatory proliferation in neighboring cells through the secretion of mitogenic signals (Tamori and Deng, 2014). Thus, we hypothesized that activation of the mitogenic AKT/FoxO signaling pathway was triggered to promote tissue repair in response to build up of tissue damage in tert−/− zebrafish. To test this hypothesis, we decided to rescue tissue degeneration by preventing p53 function and thereby unblock cell proliferation while restraining cell death. This way, we expected to thwart the induction of the AKT/FoxO proliferative pathway in older tert−/− fish and, consequently, the appearance of cell senescence.

We used tert−/− p53−/− double-mutant zebrafish where p53 deficiency rescues the adverse effects of telomere loss (Anchelin et al., 2013; Henriques et al., 2013). Consistent with our previous results, by preventing a p53-mediated response to telomere dysfunction, we were able to rescue the histopathological defects of older tert−/− testis and gut (Figure 4A and 4E). In the absence of observable tissue damage, we were no longer able to detect activation of AKT (pAKT-Ser473, Figure 4B and 4F), down-regulation of SOD2 (Figure 4B and 4F), nor accumulation of ROS (Figure 4C and 4G) in tert−/− p53−/− zebrafish. Finally, consistent with our hypothesis, older tert−/− p53−/− double-mutant gut and testis no longer exhibit an accumulation of SA-beta-Gal-positive cells (Figure 4D and 4H). Taken together, our results demonstrate that p53 is required for AKT activation and the onset of senescence in older tert−/− fish. Moreover, they suggest that the age-dependent switch from apoptosis to senescence is intimately linked to the loss of tissue homeostasis. In 3 month-old tert−/− zebrafish, telomeres are sufficiently short to trigger DDR and p53-dependent apoptosis. However, no tissue damage is observed in younger animals and this becomes apparent with age-dependent decline in cell proliferation. An older tissue with short telomeres and limited proliferative capacity responds by promoting mitogenic signalling thereby activating the AKT/FoxO pathway and consequent mitochondrial dysfunction.

### Inhibition of AKT activity prevents senescence in G1 and G2 tert−/− mutants

Our data indicates that activation of AKT in older tert−/− zebrafish correlates with the appearance of the senescence phenotype. To understand the direct role of the AKT/FoxO pathway in modulating the p15/p16-mediated cell-cycle arrest, we decided to test if AKT activation was causal to cell senescence. Our hypothesis would dictate that AKT phosphorylation inhibition would prevent p15/p16 expression and preserve tissue homeostasis.

AKT phosphorylation is mediated by the mTORC2 complex, which the main component is the mTOR (mammalian Target Of Rapamycin) protein (Laplante and Sabatini, 2009). To analyze the role of AKT activation in inducing senescence upon telomere shortening, we created a double mutant bearing a mutation in the tert gene combined with a mutation in the mTOR zebrafish homologue (zTOR). Previous work showed that zTOR is essential for development and zTOR −/− zebrafish are larval lethal (Ding et al., 2011). However, zTOR+/− mutants are haploinsufficient, with the lack of one functional copy being sufficient to reduce AKT phosphorylation (Ding et al., 2011). Thus, we decided to test our hypothesis in tert−/− ztor+/− mutant zebrafish. As expected, tert−/− ztor+/− present reduction in AKT phosphorylation compared to tert−/− single mutants in 11 month-old fish (Figure 5). Consistent with our hypothesis, this reduction is associated with a reduction in the expression of p15/16 (Figure 5), suggesting that preventing the activation of AKT can be sufficient to reduce aging-associated senescence (Figure 5 A-B). Given the incomplete nature of zTOR inhibition, haploinsufficiency for ztor in a tert−/− mutant background was insufficient to restore tissue morphology in testis and tert−/− mutant gut defects (Supp. Figure 7). Our data corroborates previous reports that disruption of zTOR partially inhibits AKT activation and, consequently, reduces p15/16 expression and with an amelioration tissue morphology of older tert−/− mutant.

Given the previous incomplete AKT inhibition in older tert−/− zebrafish, we decided to attempt a chemical inhibition in tert−/− fish with very short telomeres. Second generation telomerase-deficient zebrafish (G2 tert−/−), obtained from incross of homozygous tert mutants, recapitulate most of the phenotypes of old tert−/− G1 fish (Anchelin et al., 2013; Henriques et al., 2013). G2 tert−/− have morphological defects, along with extremely short telomeres and lifespan. Consistent with the phenotypical recapitulation of older G1 tert−/− mutants, we observed that G2 tert−/− exhibited a marked increase of senescence revealed by SA-beta-Gal staining and expression of senescence associated markers, p15/p16 (Figure 5A-B). Similar to older G1 tert−/− zebrafish, analysis of G2 tert−/− larvae showed that increase of senescence by p15/16 expression is concomitant with increased pAKT phosphorylation, decreased SOD2 and, consequently increase of ROS species (Figure 5C-D). Our data thus indicates that G2 tert−/− larvae recapitulates aging-associated AKT activation and senescence observed in old telomerase-deficient fish.

We decided to use the G2 tert−/− model to assess a direct link between AKT activation and increase in cell senescence, by testing whether direct AKT kinase inhibition would be sufficient to prevent p16 expression. For this purpose, we daily treated tert−/− and WT larvae with an AKT inhibitor (AKT 1/2 kinase inhibitor, Santa Cruz) for 2 days (Figure 5E). At 6 dpf, larvae were collected and analyzed for AKT activation and expression of senescence markers. Treatment with the inhibitor reduces AKT phosphorylation and, as a consequence, leads to a decrease of p16 protein and mRNA levels compared to controls (Figure 5F-G). Our data thus show a direct link between AKT kinase activation and senescence in tert−/− zebrafish. Taken together our results indicate that the telomere shortening-associated premature senescence is dependent on the activation of the AKT/FoxO pathway.

## DISCUSSION

Homeostasis in multicellular organisms depends on coordinated responses to external and internal insults challenging lifetime tissue integrity. Loss of tissue homeostasis is a hallmark of aging, resulting in pathologies often caused by defective or deregulated tissue damage responses (Neves et al., 2015). In proliferative tissues, homeostasis relies on a controlled balance between cell proliferation and apoptosis or senescence. Telomere attrition and DNA damage are major factors contributing to aging (López-Otín et al., 2013). When reaching a critical length, short telomeres trigger DNA Damage Responses and p53-dependent cell cycle arrest, eventually culminating in apoptosis or replicative senescence (Blackburn and Francisco, 2001; Harley et al., 1990; Olovnikov, 1973; Shay and Wright, 2000). Most cell types seem to be capable of both cellular outcomes upon damage (Campisi and d’Adda di Fagagna, 2007), but the molecular mechanism determining cell fate between apoptosis and senescence in an organism remains unclear.

In the present study, we describe that young (3 month-old) telomerase deficient zebrafish already exhibit active DDRs and p53 activation. At this stage, apoptosis is the predominant cell fate. Even though DNA damage is present in proliferative tissues, such as gut and testis, no signs of cell senescence could be detected. However, we observed a switch between apoptosis and senescence in older tert−/− fish. In these animals, senescence becomes the most prevalent cellular response, exhibiting SA-beta-Gal and p15/p16 positive cells and elevated p15/p16 and p21 levels. This observation underscores the fact that the same tissue can undergo different cellular fates, apoptosis or senescence, depending on the animal’s age.

The p53 transcription factor is described as a “master regulator” of several cellular processes, including cell cycle arrest, apoptosis, senescence and autophagy (Farnebo et al., 2010). p53 was first shown to trigger apoptosis in response to cellular stress (Vogelstein et al., 2000). However, it is now acknowledged that p53 modulates genes involved in senescence depending on the stress inflicted or cell type (Murray-Zmijewski et al., 2008). In late generation telomerase knockout mice, p53 was shown to be responsible for down-regulating of PGC1α and PGC1β and mitochondrial dysfunction upon telomere shortening (Sahin et al., 2011). In our study, early p53 activation in tert−/− zebrafish does not visibly alter mitochondria function. However, both gut and testis of older tert−/− zebrafish show mitochondrial dysfunction accompanied by significant reduction of ATP levels and accumulation of ROS. These alterations are concomitant with the onset of cell senescence. However, in contrast to the previous study in mice, we do not detect a downregulation of PGC1α either on mRNA or protein level. Even though p53 is required for the older tert−/− zebrafish phenotypes, our results suggest that the observed mitochondrial dysfunctions are independent of PGC1α alterations.

The present study reveals that mitochondrial defects are associated with a reduction in SOD2 expression allied to an Akt-dependant FoxO1 and FoxO4 phosphorylation. The anti-proliferative p53 and pro-survival mTOR/Akt pathways interact in a complex manner. Depending on the context, the interaction of these pathways modulate cell fate into either cell-cycle arrest, apoptosis or senescence (Erol, 2011; Hasty et al., 2013). Cell line studies show that p53 itself can inhibit mTOR/Akt pathway through several mechanisms including AMPK and PTEN activation (Hasty et al., 2013). Moreover, p53 activation of cell senescence relies on mTOR/Akt pathway activity (Davaadelger et al., 2016; Jung et al., 2019; Kim et al., 2017; Miyauchi et al., 2004; Vétillard et al., 2015). Consistently, Akt inhibition reduces p53-dependent senescence (Davaadelger et al., 2016; Duan and Maki, 2017; Kim et al., 2017). In addition, Akt mediates the inhibition of pro-apoptotic factors (Davaadelger et al., 2016) and leads to increased levels of anti-apoptotic Bcl-xl (Jones et al., 2000; X. Li et al., 2017). Thus, activation of mTOR/Akt pathway can act as a negative regulator of apoptosis (Davaadelger et al., 2016; Duan and Maki, 2017). Akt was shown to induce senescence and cell-cycle arrest by elevating the intracellular levels of ROS or activating p16 transcription through direct phosphorylation of the Bmi repressor (Imai et al., 2014; L. U. Li et al., 2017; Liu et al., 2012; Miyauchi et al., 2004; Nogueira et al., 2008). Accordingly, Akt activation in aged tert mutant zebrafish is concomitant with increased anti-apoptotic Bcl-xl and pro-senescence p15/p16 and p21 levels.

What constitutes the mechanistic nature of the switch from apoptosis to senescence? Even though young tert−/− mutants present no observable tissue defects, they exhibit high levels of apoptosis and a reduction in proliferative capacity (Carneiro et al., 2016a; Henriques et al., 2013). High apoptosis increases the demand of cell proliferation from surrounding cells in a process termed apoptosis-induced compensatory proliferation (Fan and Bergmann, 2008). Thus, tissue degeneration becomes apparent in aging tert−/− zebrafish. In tissues where stem cells are not readily available or where tissue-intrinsic genetic programs constrain cell division, cellular hypertrophy represents an alternative strategy for tissue homeostasis (Losick et al., 2013; Tamori and Deng, 2013).

We propose that, upon telomere shortening and p53 activation, loss of tissue integrity triggers the AKT-dependent pro-proliferative pathway (Figure 3). The combination of these antagonistic forces in the cell would result in cellular senescence. We tested this hypothesis on both pathways. By genetically removing tp53, we were able to rescue tissue degeneration and avoid activation of AKT, accumulation of ROS and induction of senescence. On a second level, we inhibited TOR/Akt genetically by dampening the ztor pathway and, chemically, by directly inhibiting Akt in G2 tert−/− larvae. In both cases, we were able to reduce the effects of telomere shortening. Collectively, our results show that the crosstalk between two pathways telomere shortening/DDR and AKT/FoxO signalling regulate a apoptosis-to-senescence switch and contributes to tissue homeostasis *in vivo*.

## Acknowledgements

We thank members of the Telomeres and Genome Stability Laboratory for helpful discussions. We thank the Instituto Gulbenkian de Ciência EM unit, histology unit and the Fish Facility for excellent animal care. MEM is a recipient of a postdoctoral fellowship from the Ville de Nice. This work was supported by the FCT (PTDC/SAU-ORG/116826/2010), Fondation Arc pour la Recherche sur le Cancer (PJA20161205137) and the Howard Hughes Medical Institute International Early Career Scientist grants received by MGF.

## Author contributions

Conceived and designed the experiments: MGF, IC, MEM and MM. Performed the experiments: MEM, MM and IC. Analysed the data: IC, MEM, MM and MGF. Contributed reagents/materials/analysis tools: IC, MEM and MM. Wrote the paper: MGF, MEM, MM and IC.

## MATERIALS AND METHODS

### Ethics statement

All Zebrafish work was conducted according to National Guidelines and approved by the Ethical Committee of the Instituto Gulbenkian de Ciência and the DGAV (Direcção Geral de Alimentação e Veterinária, Portuguese Veterinary Authority).

### Zebrafish lines and maintenance

Zebrafish were maintained in accordance with Institutional and National animal care protocols. The telomerase mutant line tertAB/hu3430, generated by N-Ethyl-N-nitrosourea (ENU) mutagenesis (Utrecht University, Netherlands; Wienholds, 2004), has a T-A point-mutation in the tert gene. tertAB/hu3430 line is available at the ZFIN repository, ZFIN ID: ZDB-GENO-100412-50, from the Zebrafish International Re-source Center—ZIRC. The protocols used for outcrossing mutagenized male zebrafish were previously described (Carneiro et al., 2016b; Henriques et al., 2013). The terthu3430/hu3430 homozygous mutant (tert−/−) was obtained by incrossing our tertAB/hu3430 strain. WT siblings were used as controls. Overall characterization of tert−/− and WT zebrafish was performed in F1 animals produced by tert+/− incross. Due to a male sex bias in our crosses that affected mostly tert−/− progeny, we were unable to obtain significant numbers of females for analysis and so all of our data is restricted to males.

p53 mutant line zdf1 (P53M214K) (Berghmans et al., 2005) was kindly provided by António Jacinto (CEDOC, chronic diseases, Nova medical school, Lisbon (Portugal)). p53 mutant line zdf1 (P53M214K) is available at the ZFIN repository, ZFIN ID: ZDB-ALT-050428-2 from the Zebrafish International Re-source Center—ZIRC. The p53M214K/ p53M214K homozygous mutant (p53−/−) was obtained by incrossing our p53AB/M214K strain.

ztor line was obtained from the ZFIN repository, ZFIN ID: ZDB-ALT-120412-1from the Zebrafish International Re-source Center—ZIRC. The line was previously described (Ding et al., 2011) as homozygous larval lethal and it was maintained through outcrossing. All animals showing signs of morbidity that persisted for up to 5 days, such as inability to eat or swim, or macroscopic lesions/tumors were sacrificed in 200 mg/L of MS-222 (Sigma,MO, USA).

### Real-time quantitative PCR

Age- and sex-matched fish were sacrificed in 200 mg/L of MS-222 (Sigma, MO, USA) and portions of each tissue (gonads, gut and muscle) were retrieved and immediately snap-frozen in liquid nitrogen. Similarly, 4dpf larvae were sacrificed and collected in Eppendorf tube, minimum 10 larvae each. RNA extraction was performed in TRIzol (Invitrogen, UK) by mashing each individual tissue with a pestle in a 1.5 ml eppendorf tube. After incubation at RT for 10 minutes in TRIzol, chlorophorm extractions were performed. Quality of RNA samples was assessed through BioAnalyzer (Agilent 2100, CA, USA). Retro-transcription into cDNA was performed using a RT-PCR kit NZY First-Strand cDNA Synthesis Kit # MB12501 (NZYtech).

Quantitative PCR (qPCR) was performed using iTaq Universal SYBR Green Supermix # 1725125 (Bio-Rad) and an ABI-QuantStudio 384 Sequence Detection System (Applied Biosystems, CA, USA). qPCRs were carried out in triplicate for each cDNA sample. Relative mRNA expression was normalized to rpl13 a (data not shown) mRNA expression using the DCT method. Primer sequences are listed in Table S1.

**Table S1.**
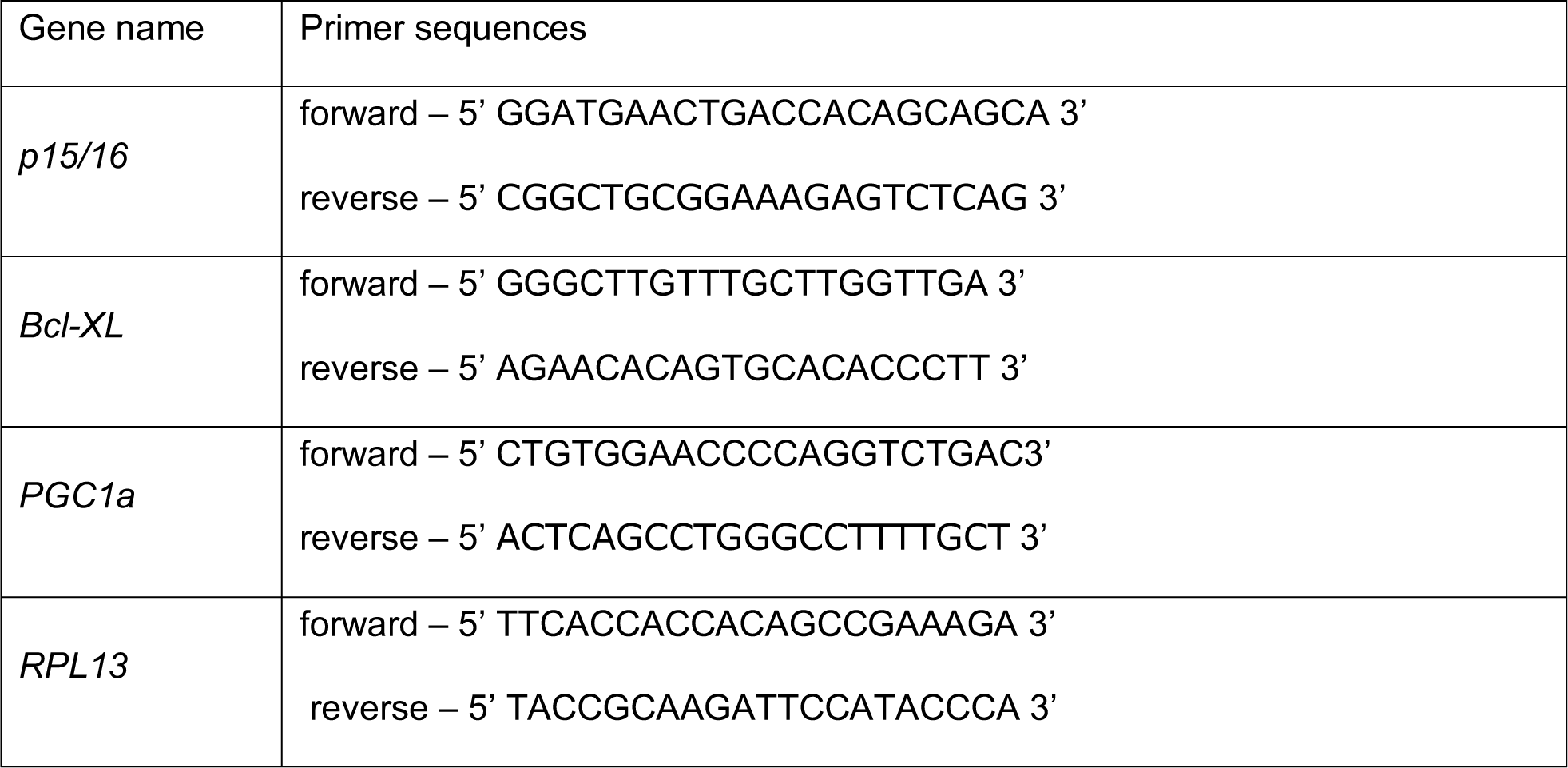
List of primers used in RT-qPCR expression analysis and *tert* genotyping.

### Detection of intracellular oxidant activity

Reactive oxygen species (ROS) accumulation was assessed by measuring the levels of the oxidized form of the cell-permeant 5-chloromethyl-2’,7’-dichlorodihydrofluorescein diacetate (DCFDA) (Sigma). Briefly, zebrafish were euthanized with 200 mg/L of MS-222 (Sigma, MO, USA) and tissues such as the testis, gut and muscle were dissected. Each tissue was homogenized in 100 μl of ROS buffer (0.32 mM sucrose, 20mM hepes, 1mM MgCl2 and 0.5mM Phenylmethanesulphonyl fluoride). Homogenates were centrifuged and 20 μl of the supernatant was transferred to a 96-well plate and incubated in 1 μg/ml of DCFDA for 30 minutes. Fluorescence values were measured with a Victor 3 plate reader (Perkin Elmer) and normalized to total protein content, which was determined by the Bradford method. N = 3 per time point.

### Histological analysis

Zebrafish were sacrificed by anaesthetic overdose, in 200 mg/L of MS-222 (Sigma, MO, USA), fixed for 72 hours in 10% neutral buffered formalin and decalcified in 0.5MEDTA for 48 h. Whole fish were then paraffin-embedded and 3 micrometer midline sagittal sections were stained with haematoxylin and eosin for histopathological analysis. Sections were examined by a pathologist (TC), blinded to experimental groups and microphotographs were acquired in a Leica DM2500 microscope coupled to a Leica MC170 HD microscope camera. At least 4 animals from each age group/genotype were analysed.

### Immunofluorescence (IF) and confocal analysis

Apoptosis and Senescence was detected using the In Situ Cell Death Detection Kit (Roche, SW) according to manufacturer’s instructions combined with Immunofluorescence against the p15/16 senescence-associated factor. Briefly, deparaffinized slides were incubated with 40 μg/ml Proteinase K in 10 mM Tris-HCl pH 7.4, 45 minutes at 37°C. Slides were left to cool down for 30 minutes at room temperature (RT), washed three times in dH20 for 5 minutes each and blocked for 1 hour at RT in 1% BSA, 0,5% Tween 20 in PBST (Triton 0.5%). Subsequently the slides were incubated over-night with anti-p16 (F-12) (1:50, Santa Cruz Biotechnology, sc-1661), followed by 3×10 minute PBS washes. Incubation with the secondary antibody Alexa Fluor 568 goat anti-mouse (Invitrogen, UK, 1:500 dilution) overnight at 4°C was followed by three 10 minute PBS washes. The day after the slides were washed 2×5 minutes in PBS and then incubated with TUNEL labelling mix (protocol indicated by the supplier). Washing and mounting were performed by DAPI staining (Sigma, MO, USA) and mounting with DAKO Fluorescence Mounting Medium (Sigma, MO, USA).

Images were acquired on a commercial Nikon High Content Screening microscope, based on Nikon Ti equipped with a Andor Zyla 4.2 sCMOS camera, using the a 20x 1.45 NA objective, DAPI + GFP fluorescence filter sets and controlled with the Nikon Elements software.

For quantitative and comparative imaging, equivalent image acquisition parameters were used. The percentage of positive nuclei was determined by counting a total of 500–1000 cells per slide, 63x amplification (N = 3–4 zebrafish per time point/genotype).

### Senescence-associated β-galactosidase assay

β-galactosidase assay was performed as previously described (Kishi et al., 2008). Briefly, sacrificed zebrafish adults were fixed for 72h in 4% paraformaldehyde in PBS at 4°C and then washed three times for 1 h in PBS-pH 7.4 and for a further 1 h in PBS-pH 6.0 at 4°C. β-galactosidase staining was performed for 24 h at 37°C in 5 mM potassium ferrocyanide, 5 mM potassium ferricyanide, 2mM MgCl2 and 1 mg/ml X-gal, in PBS adjusted to pH 6.0. After staining, fish were washed three times for 5 minutes in PBS pH 7 and processed for decalcification and paraffin embedding as before. Sections were stained with nuclear fast red for nuclear detection and images were acquired in a bright field scan (Leyca, APERIO).

### Statistical and image analysis

Image edition was performed using Adobe Photoshop CS5.1 Statistical analysis was performed in GraphPad Prism5, using two-way ANOVA test with Bonferroni post-correction for all experiments comparing WT and tert−/− over time. For real-time quantitative PCR, statistical analysis was performed in GraphPad Prism5, two-way ANOVA with Bonferroni post-correction. A critical value for significance of p<0.05 was used throughout the study. For Western Blot the bands intensities were calculated using FIJI. Statistical analysis was performed using GraphPad Prism6, the significance was assigned according to the Mann-Whitney t-test. A critical value for significance of p<0.05 was used throughout the study.

### Immunoblot analysis

Age- and sex-matched adult zebrafish fish were sacrificed in 200 mg/L of MS-222 (Sigma, MO, USA) and portions of each tissue (gonads and gut) were retrieved and immediately snap-frozen in dry ice. 4dpf larvae were sacrificed in ice and collected in 1,5mL Eppendorf tube, minimum 10 larvae /tube. Gonads tissues and larvae were then homogenized in RIPA buffer (sodium chloride 150mM; Triton-X-100 1%; sodium deoxycholate 0,5%; SDS 0,1%; Tris 50mM, pH=8.0), including complete protease and phosphatase inhibitor cocktails (Roche diagnostics), with the help of a motor pestle. Protein extracts were incubated on ice for 30 minutes and centrifuged at 4ºC, 13.000 rpm, for 10 min. The supernatant was collected and added to 100 mL of protein sample buffer containing DTT.

Gut samples were homogenized in TRIzol (Invitrogen, UK) by mashing each individual tissue with a pestle in a 1.5 ml Eppendorf tube. After incubation at RT for 10 minutes in TRIzol, chlorophorm extractions were performed. The organic phase was collected and proteins were precipitated according to the manufacture protocol. The protein pellet was resuspended in 100ul of Lysis Buffer (150mM NaCl, 4%SDS, 50mM TrisHCl pH 8.0, 10mM EDTA, complete protease and phosphatase inhibitor cocktails-Roche diagnostics).

For each sample, a fraction of Proteins was separated on 12% SDS-PAGE gels and transferred to Immobilon PVDF membranes (Millipore). The membranes were blocked in 5% milk or 5% BSA (depending on the primary antibody), then incubated with the indicated primary antibody prior to incubation with the appropriate HRP-conjugated secondary antibody. Antibody complexes were visualised by enhanced chemiluminescence (ECL). Antibodies concentration: anti-Tp53 (1:1000, Anaspec, 55342); anti-g-H2AX (1:1000, GeneTex, GTX127342); anti-p16 (F-12) (1:700, Santa Cruz Biotechnology, sc-1661); anti-SOD-2 (1:1000, Sigma, SAB2701618); anti-phospho-AKT, Ser473 (1:1000, Cell Signaling, #4060); anti-total-Akt (1:1000 Cell Signaling, #9282, gift of Adrien Colin), anti-phospho-FoxO1, Ser256 (1:100, Cell Signaling, #9461); anti-Tubulin (1:5000, Sigma, T 6074).

### ATP measurement

Age- and sex-matched adult zebrafish fish were sacrificed in 200 mg/L of MS-222 (Sigma, MO, USA) and portions of each tissue (gonads, gut and muscle) were retrieved and immediately snap-frozen in dry ice. Each tissue was homogenised in 100 mL of 6M guanidine-HCl in extraction buffer (100mM Tris and 4mM EDTA, pH 7.8) to inhibit ATPases. Homogenised samples were subjected to rapid freezing in liquid nitrogen followed by boiling for 3 minutes. Samples were then cleared by centrifugation and the supernatant was diluted (1/50) with extraction buffer and mixed with luminescent solution (CellTiter-Glo Luminescent Cell Viability Assay, Promega). The luminescence was measured on a Victor 3 plate reader (Perkin Elmer). The relative ATP levels were calculated by dividing the luminescence by the total protein concentration, which was determined by the Bradford method. For Bradford assays, samples were diluted (1/50) with extraction buffer.

### Electron microscopy

For electron microscopy analysis, zebrafish tissues were processed according to Schieber et al, 2010. Briefly, zebrafish were fixed in 2% Paraformaldehyde, 2.5% Glutaraldehyde in 0.1M PHEM buffer for 72h at 4°C. Dissected tissues were then washed 3 times in 0.1M PHEM. Tissues were transferred in 1% Osmium Tetroxide in 0.1M PHEM for 1h fixation on ice. Samples were then dehydrated before being processed for embedding using Epon (Schieber et al., 2010). 70 nm ultrathin sections were cut using Reichert Ultramicrotome. After being counterstained with uranyl acetate and lead, samples were analyzed using a transmission electron microscope (Hitachi H-7650).

### AKT inhibitor larval treatment

AKT ½ kinase inhibitor (AKT inh) was purchased from Santa-Cruz (sc-300173). Stock solutions were prepared in DMSO. AKT inh was applied, after a titration, at 2uM concentration between days 3 and 5 post fertilization. Larvae were grown at 28°C and over the incubation periods, replacement of medium with the above mentioned compounds was performed every day, between 3 and 7 PM. Since the compound was dissolved in DMSO, controls were treated with the correspondent dilution of the solvent. The drug was tested in 2 independent trials. Finally, 5dpf larvae were sacrificed and collected to perform protein and RNA analysis.

### Gene knock-down using p15/16 Morpholino injection

One-cell stage WT embryos were injected with 2.4 ng or 3.6 ng of p15/16 mRNA specific translation blocking antisense morpholino oligonucleotides (MO) sequence (5’ TCAGTTCATCCTCGACGTTCATCAT 3’) or 3.6 ng of standard control MO (5’ CCTCTTACCTCAGTTACAATTTATA 3’) (Gene Tools, USA). After 4 dpf, larvae were collected for further p15/p16 protein expression analysis.

---

**Supplemental Figure 1.**
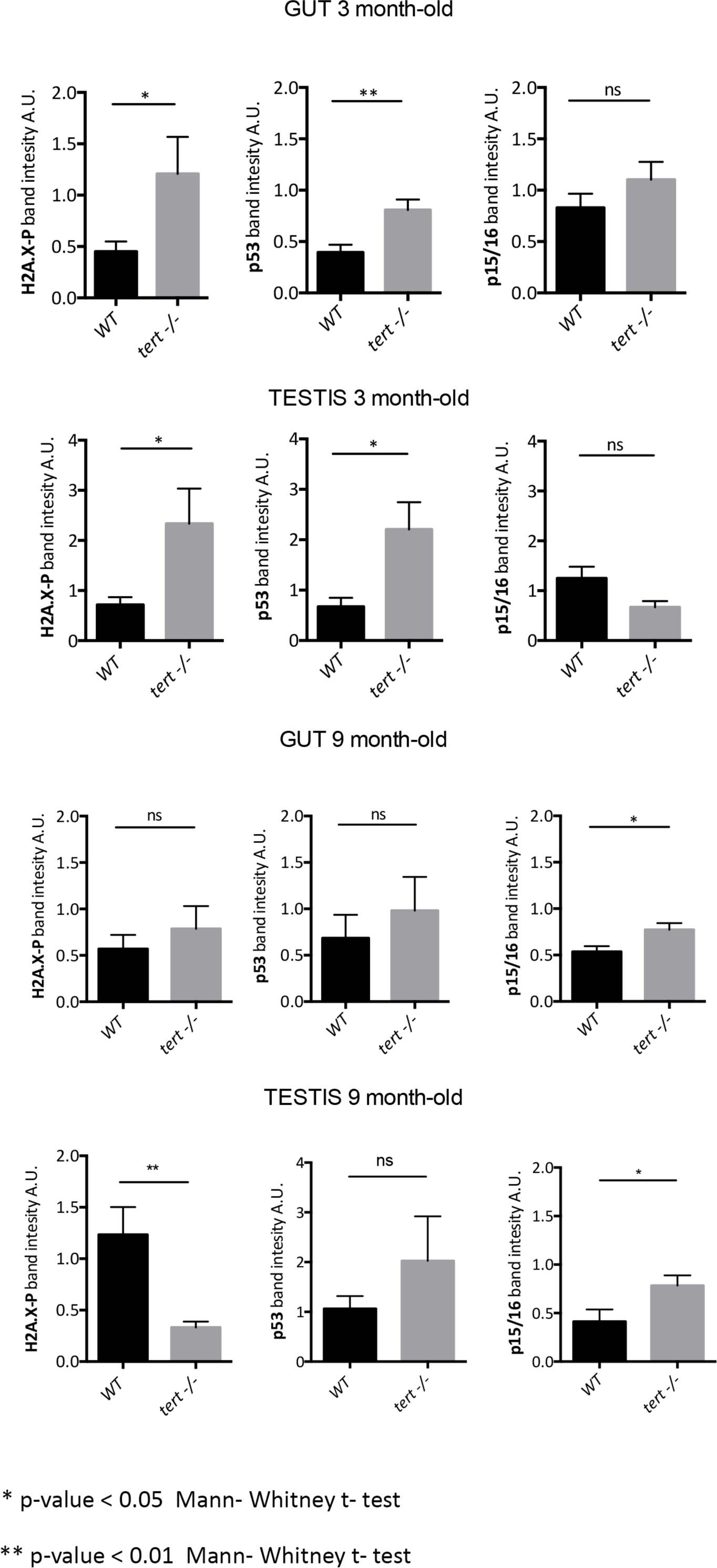
Proliferative tissues of tert−/− zebrafish exhibit a switch from apoptosis to senescence with age. Quantification of Western blot analysis for DNA Damage and senescence associated genes (depicted in Fig1) in gut and testis of 3 month or 9 month-old WT and tert−/− siblings (N>=6 fish). At 3 month, gut and testis showed higher levels gH2AX and p53 proteins in tert−/− compared to WT but no differences in P15/p16 expression. At 9 month, senescence-associated p15/16 expression increased in tert mutant compared to WT siblings. Band intensities were analysed by ImageJ and normalised by Tubulin or Actin control bands. Data are represented as mean±SEM. * p-value <0.05; ** p-value<0.01 using Mann-Whitney t-test.

**Supplemental Figure 2.**
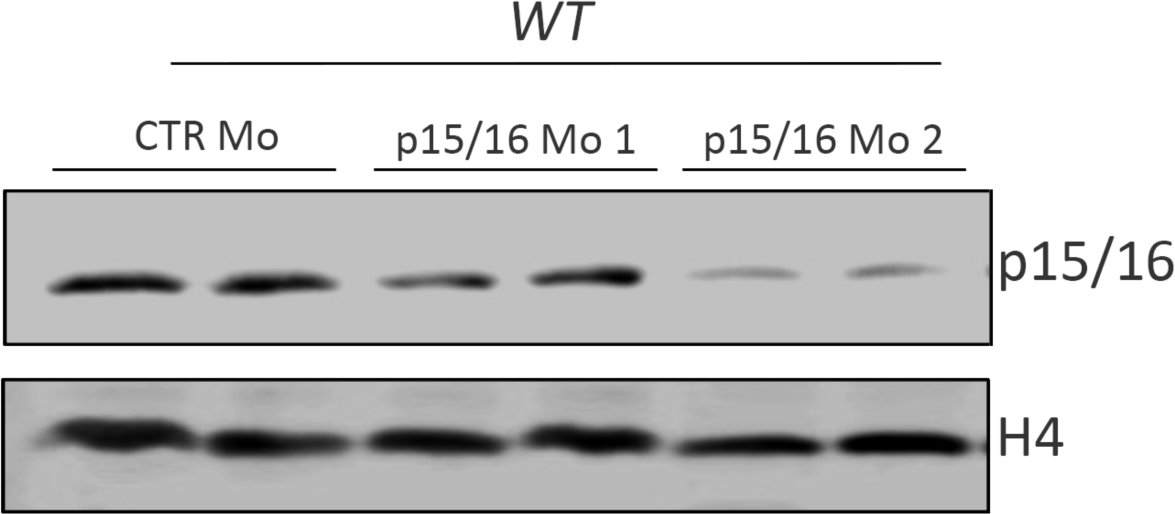
Anti-p16 antibody validation in zebrafish through antisens morpholino oligonucleotide knock-down of p15/p16. Representative Western blot of p15/p16 using pool of 4dpf larvae injected (at 1 cell-stage) with control, 2.4 ng or 3.6 ng of p15/p16 antisens morpholino oligonucleotides. Dose-dependent decrease of p15/p16 protein levels with p15/p16 morpholinos confirm the specificity of anti-p16 (F-12) (1:50, Santa Cruz Biotechnology, sc-1661) for zebrafish p15/p16 protein.

**Supplemental Figure 3.**
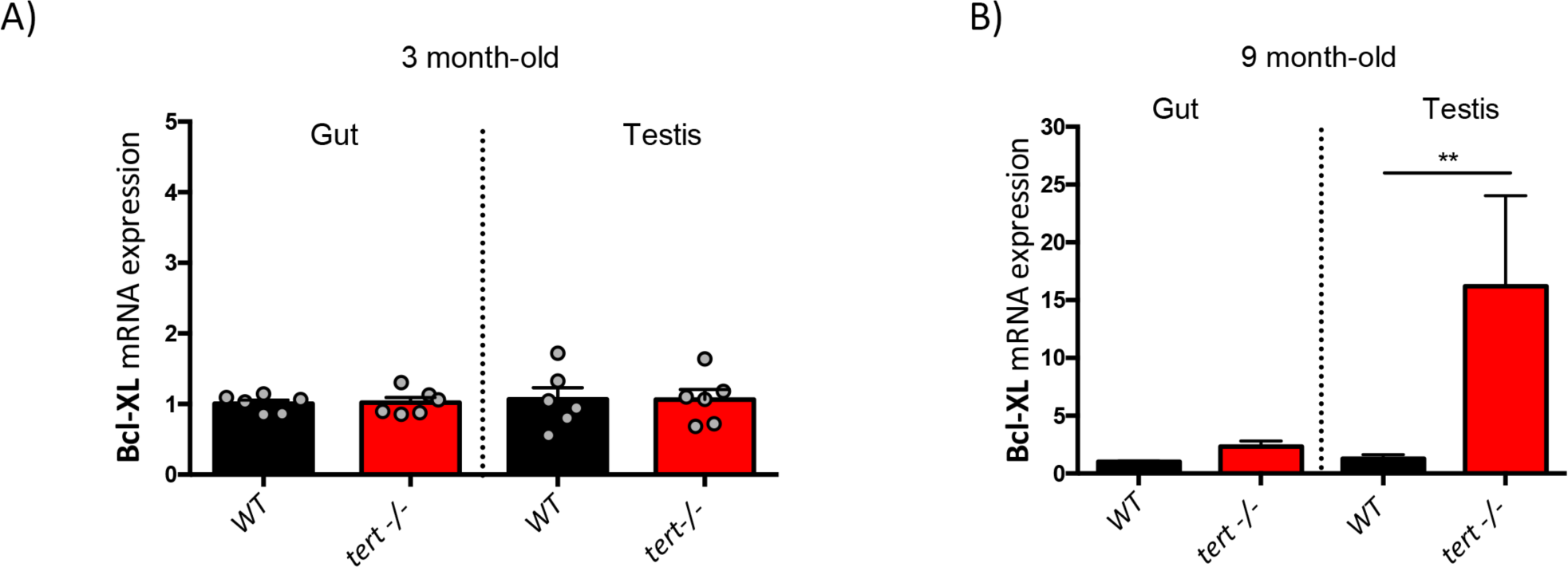
Bcl-XL is overexpressed in 9month but not 3month-old tert−/−. RT-qPCR analysis of Bcl-XL in gut and testis of 3 or 9 month-old tert−/− or WT siblings (N=6 fish). The graphs are representing mean ±SEM mRNA fold increase after normalisation by RPL13a gene expression levels (* p-value <0.05; ** p-value<0.01). While no differences are seen at 3 months, Bcl-XL is overexpressed in 9 month-old tert−/− gut and testis compared to WT.

**Supplemental Figure 4.**
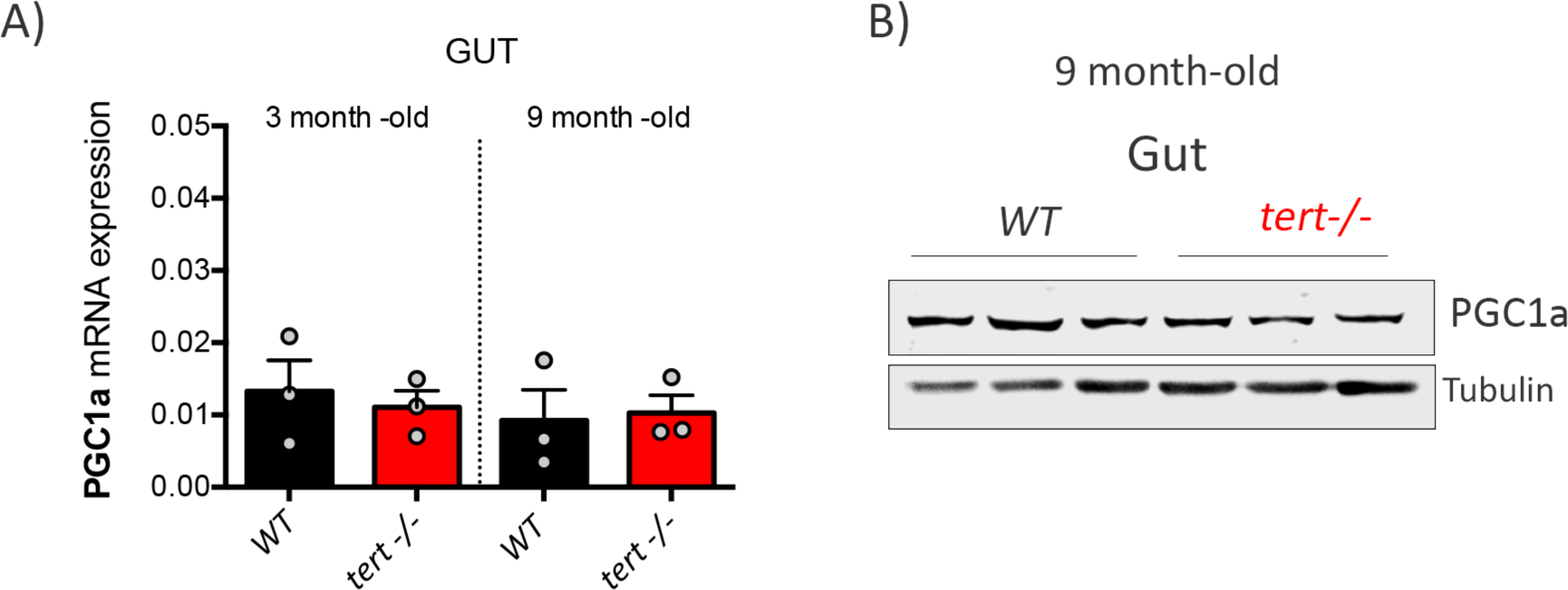
PGC1a expression is not altered in tert mutants compared to WT. A) RT-qPCR and B) representative Western blot analysis of PGC1a in gut of 3 or 9 month-old tert−/− or WT siblings (N=3 fish). The graphs are representing mean ±SEM mRNA fold increase after normalisation by RPL13a gene expression levels (* p-value <0.05; ** p-value<0.01). No differences are detected in PGC1a expression between tert−/− and WT gut at 3 and 9 month.

**Supplemental Figure 5.**
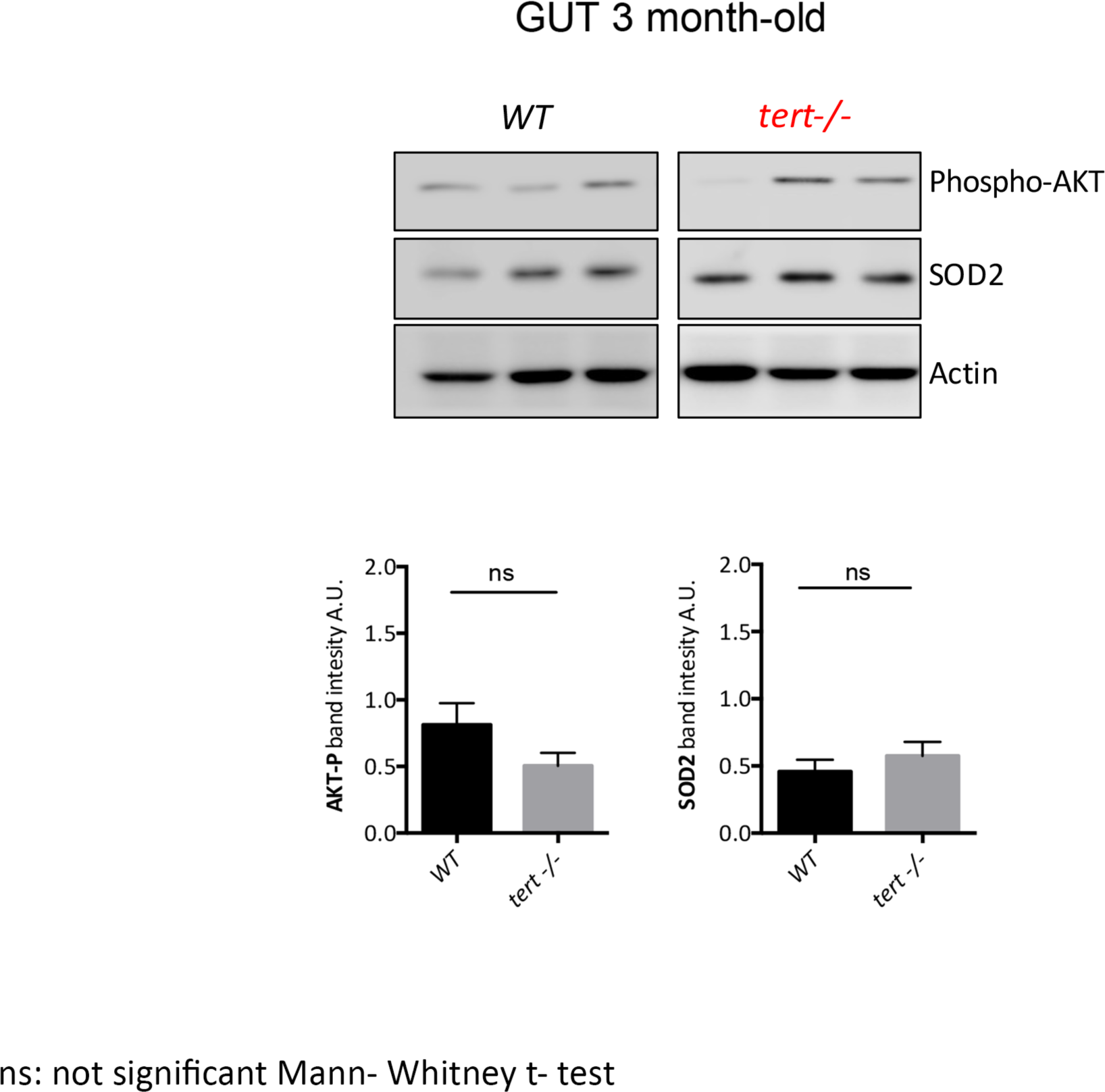
Akt pathway is not activated in young tert−/− compared to wild-type. Representative immunoblot of p-Akt and SOD2 and related quantification from testis and gut of 3 month-old tert mutant and WT siblings (N=3). At 3 month, no differences are observed in p-Akt and SOD2 protein levels between tert−/− and WT siblings. Data are represented as mean±SEM.

**Supplemental Figure 6.**
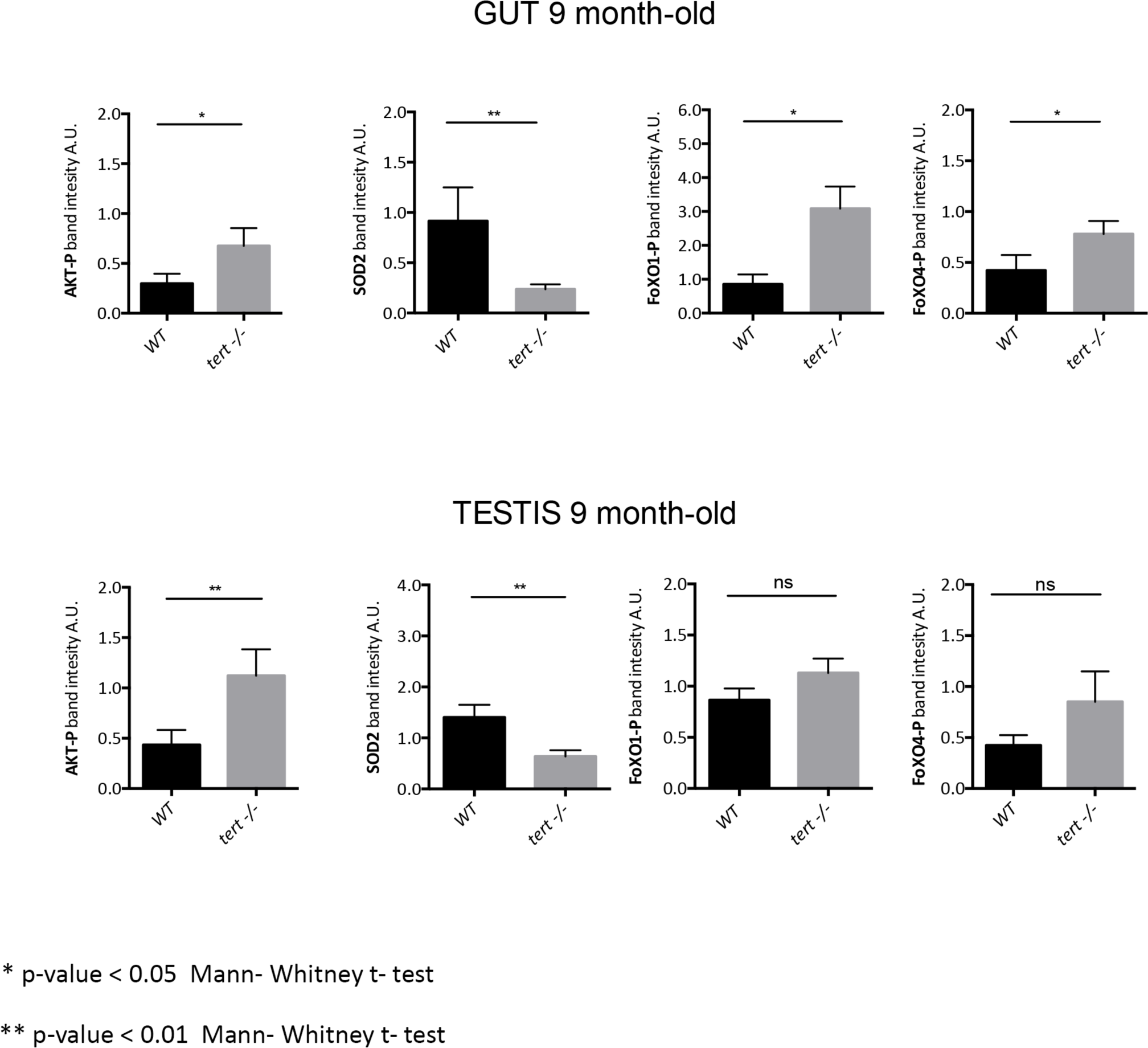
Activation of Akt in old tert−/− leads to ROS accumulation by blocking the FoxO1/4-SOD2 Axis and promoting mitochondrial dysfunction. Quantification of Western blot of p-Akt, pFOXO1, pFOXO4 and SOD2 (depicted in Fig3) from testis and gut of 9 month-old tert mutant and WT siblings (N>=9). At 9 month, these proliferative tissues show an increased activation of Akt leading to the inhibition of FOXO-dependant SOD2 expression. Band intensities were analysed by ImageJ and normalised by Tubulin control bands. Data are represented as mean±SEM. * p-value <0.05; ** p-value<0.01 using Mann-Whitney t-test.

**Supplemental Figure 7.**
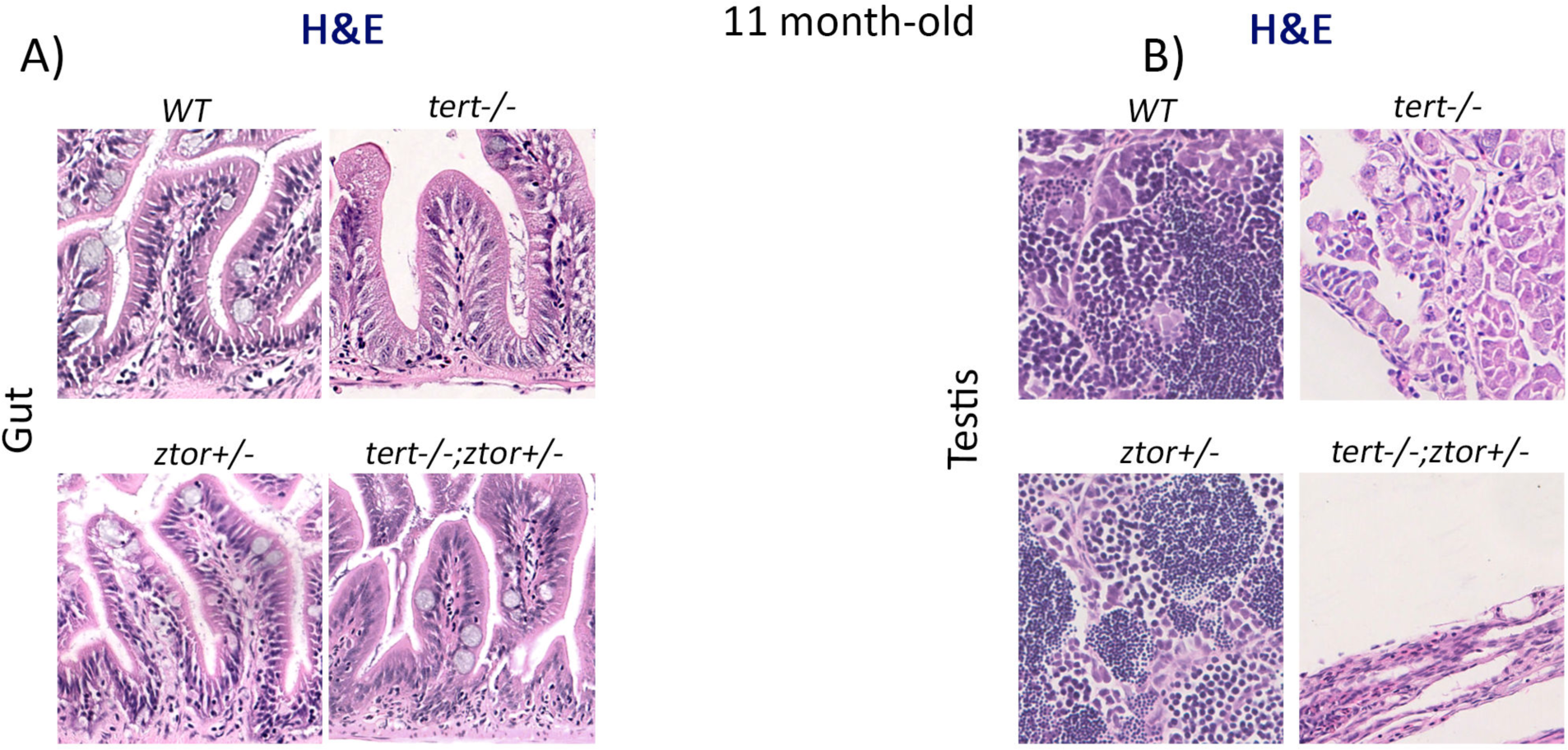
ztor haploinsufficiency is not sufficient to prevent tissue defects in tert−/− zebrafish. A and B) Representative haematoxylin and eosin-stained sections of gut (A) and testis (B) from 11 month-old WT, tert−/−, ztor+/− and tert−/−; ztor +/− siblings (N=3 fish each). The absence of one copy of the ztor gene is not sufficient to rescue the morphological defects observed in the tert −/− at 11 month of age.

